# Liposome-assisted in-situ cargo delivery to artificial cells and cellular subcompartments

**DOI:** 10.1101/2022.04.27.489538

**Authors:** Lin Xue, Anna B. Stephenson, Irep Gözen

## Abstract

We report on liposome-mediated targeted delivery of membrane-impermeable constituents into surface-adhered giant lipid compartments, employed as artificial cells. Soluble cargo compounds are delivered by means of an open-space microfluidic device, which perfuses selected lipid compartments with loaded small unilamellar vesicles (SUVs) composed of cationic lipids. The SUV membranes fuse with the surface-adhered containers, merging their contents. We monitored the fusion process via Förster resonance energy transfer (FRET) by labeling both the membranes of the SUVs and the target compartments with a fluorophore pair. We established that, upon fusion, water-soluble dyes, fluorescently labeled genetic polymers, sugars and proteins carried by the SUVs can be successfully internalized at high yield. Finally, by transferring carbonic anhydrase (CA) to the giant lipid compartments, enzymatic hydrolysis of the prefluorescent carboxyfluorescein diacetate (CFDA) is demonstrated by the emission intensity increase emanating from the product carboxyfluorescein (CF). Spontaneous subcompartmentalization occurred during liposomal delivery of the enzyme, leading to CF formation in an organelle-like subcompartment. The reported targeted delivery technique enables chemical reactions and cell-free gene expression in synthetic cell models with unprecedented ease and precision, and opens pathways to protocell architectures with distinct functional subcompartments in the context of origins of life research.

## Introduction

Artificial cells are simplified models of biological cells and are commonly employed in order to understand complex structural and functional features of cells^*1*^. Typically, membranous compartments, e.g. giant unilamellar vesicles (GUVs), serve as minimal cell models^*2-9*^. GUVs are comparable to biological cells in size, and they enclose an aqueous volume inside a continuous spherical phospholipid bilayer, similar to plasma membrane-enveloped contemporary biological cells. GUVs can be easily generated by self-assembly and visualized by light microscopy in real time. Towards gradually building biomimetic entities of increased complexity, GUVs were shown to accommodate protein reconstitution^*10*^ as well as cell-free gene expression^*11*^, and to undergo shape transformations such as fusion^*12*^ and division^*8, 13*^.

In order to achieve the structural and functional resemblance to living cells, a common approach is to encapsulate biological constituents inside the giant vesicles. Encapsulating macromolecules and large substructures inside the GUVs is a considerable challenge. Efficient methods exist, such as injection via glass microcapillaries^*14-16*^, pulse-jetting^*17*^ or other microfluidics based methods, but they are often cumbersome and require highly specialized instrumentation. Methods based on continuous droplet interface crossing encapsulation^*18, 19*^ and double emulsions^*20*^, are more convenient to setup and apply and are efficient in terms of encapsulation, but the solvents and oils used in such processes are difficult to completely eliminate from the resulting samples. Such residues can cause structural defects and interfere with the chemical reactions inside the artificial cells. Moreover, most techniques address bulk systems, and are not applicable to surface-adhered containers, which are preferable for microscope observation and sequential manipulation.

The use of cationic lipid particles for transfection of biological cells is a well-established approach to internalization^*21, 22*^. There have been a few studies which employ this method for delivery of cargo to GUVs, e.g. fluorescent dye calcein^*23*^ and DNA^*24*^. The mechanisms of fusion of SUVs to GUV membranes has been recently studied, but only with cargo-free liposomes^*25*^.

In this work, we utilize cationic liposomes to deliver different types of membrane-impermeable cargo: a water-soluble dye, oligonucleotides, a complex branched polysaccharide, a tetrameric protein and an enzyme, to surface-adhered GUVs in-situ. We used a microfluidic pipette for local delivery of cargo to each surface-adhered GUV, and monitored the delivery in real time with confocal microscopy. Delivery of an enzyme was achieved in conjunction with spontaneous formation of subcompartments inside a surface-adhered artificial cell, followed by the hydrolysis of the substrate and generation of fluorescence product. The SUV-mediated delivery technique we employed can be used to internalize reactants, enzymes or similar, to perform pre-or probiotic reactions and cell-free gene expression, in surface-adhered artificial cells.

## Results and Discussion

The schematic drawing of our experimental setup is shown in **Fig. 1**. In brief, GUVs (d≈10 μm, magenta color), adhered on a solid surface submerged in an aqueous solution, are exposed to cargo-loaded fusogenic SUVs, i.e. liposomes (d=100 nm, yellow color)^*25, 26*^ by using a commercially available microfluidic pipette^*27*^. The pipette creates a limited superfusion zone around each GUV and exposes it locally to the cationic liposomes^*28*^. During exposure, the SUVs fuse to the GUV membrane, resulting in delivery of the cargo molecules to the interior volume of the GUV. The GUV membrane, the SUV membrane, and the cargo molecules were individually fluorescently labeled. Both the exposure of the liposomes to the GUV and the delivery of cargo can thus be observed with an inverted laser scanning confocal microscope in real time. While vesicle fusion and cargo delivery has previously been achieved by injecting the SUVs into a GUV suspension^*24*^ or into adherent cells^*29*^ and vesicles^*23*^ using a hand-held automated pipette, an open-volume device (microfluidic pipette) with a limited exposure zone process allows focusing on one vesicle at a time, while providing accurate control over the delivered amounts of cargo.

**Figure 1.**
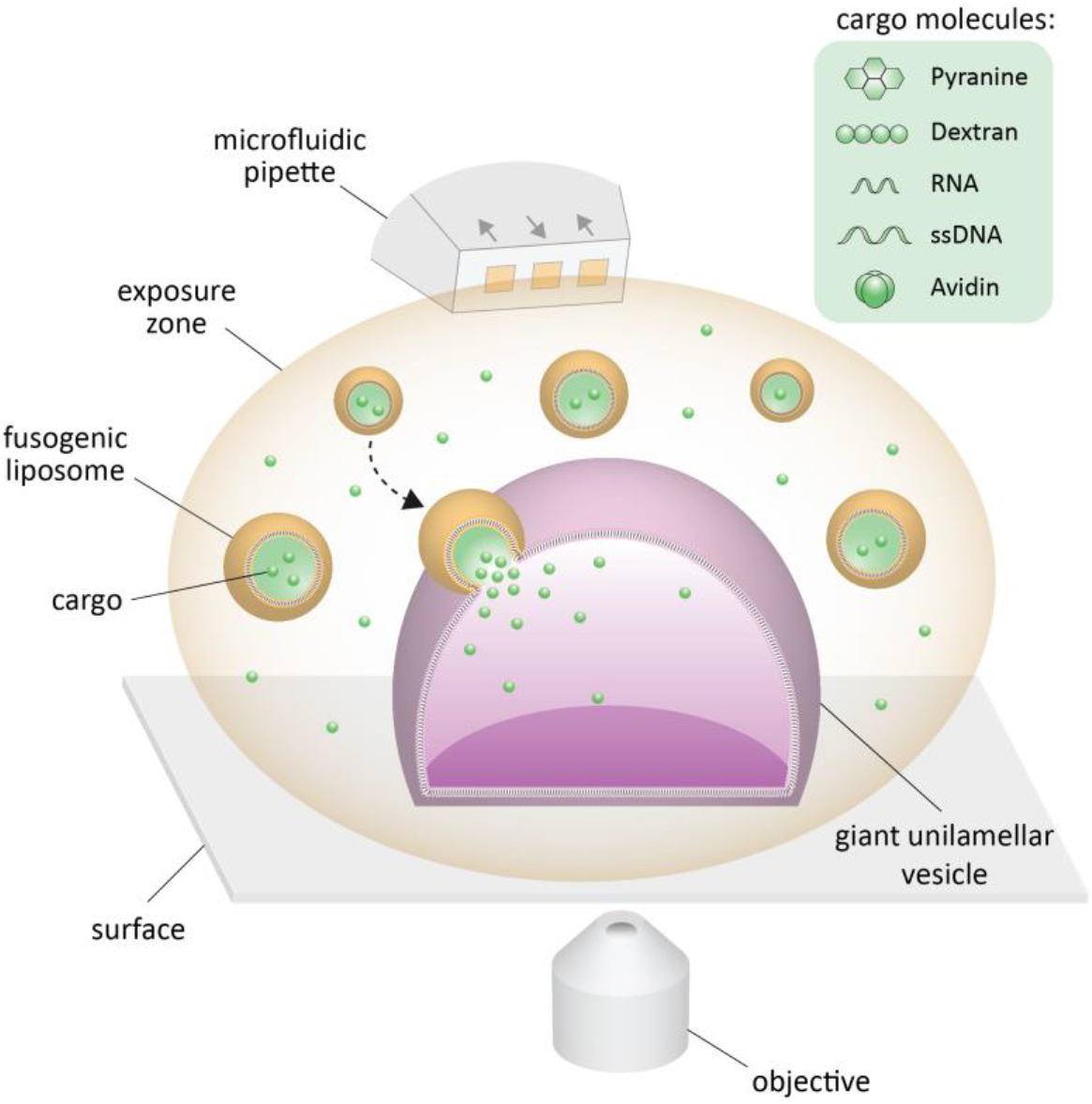
Schematic drawing of the experiment (not to scale). A surface-adhered GUV is locally exposed to fusogenic liposomes by using a microfluidic pipette. The pipette can create an exposure zone in which different cargo molecules can be re-circulated locally around the GUV, e.g. pyranine, dextran, RNA and DNA oligomer and avidin. Upon fusion of the liposomes with the GUV membrane, the initially encapsulated cargo molecules are released into the giant vesicle. All experiments have been performed in aqueous buffers.

The labeled cargo molecules used in this study are listed in **Fig. 1**: water-soluble dye pyranine, a branched polysaccharide (dextran), RNA, single-stranded (ss) DNA, and the protein avidin.

Pyranine is a low molecular weight (0.52 kDa), pH sensitive^*30*^ and water-soluble fluorescent dye which is commonly used in cells and lipid vesicles to monitor the internal pH^*31, 32*^. Polysaccharide dextran is one of the commonly used biopolymers to achieve compartmentalization inside lipid vesicles via liquid-liquid phase separation^*33, 34*^. Avidin (*m*=66 kDa) has high affinity for biotin which can be conjugated to lipid membranes or nanoparticles. Avidin-biotin interactions have been commonly employed throughout biochemical assays and functionalization of GUVs, e.g. to tether nanoparticles on GUV membranes^*35*^, immobilize actin filaments on the inner leaflet of GUV membranes^*9*^, or to induce domain formation in lipid/polymer hybrid vesicles^*36*^. GUVs are suitable primitive cell models in origin of life studies; encapsulation of genetic fragments is of particular interest in this context. The widely accepted RNA world hypothesis^*37*^, suggests that RNA could have been the component which carried genetic information, self-replicated and catalyzed chemical reactions^*38*^ prior to the emergence of DNA and proteins. Later, DNA emerged, which due to its higher stability assumedly became the predominant genetic material^*37, 39*^. In synthetic biology, RNA and DNA have been utilized to perform cell-free gene expression^*40*^ inside lipid vesicles^*11, 41*^ aiming at pathways to the bottom up construction of a living cell.

### Fusion of small fusogenic liposomes to giant vesicles

First, we performed fusion experiments using liposomes which do not carry any cargo (**Fig. 2**). Liposomes composed of positively charged lipids fuse with neutrally or negatively charged GUVs upon contact (**Fig. 2a**). We labeled the GUVs and SUVs with lipid-conjugated dyes, ATTO 488 (FRET donor) and ATTO 655 (FRET acceptor), which are suitable for FRET^*42*^. FRET has been used previously to visualize lipid mixing^*25*^. **Fig. 2b** shows the confocal 3D micrograph of a surfaced-adhered GUV before liposome delivery. Within three minutes of liposome exposure and membrane fusion, fusogenic lipids are transferred into the GUV membrane, resulting in an increased fluorescence emission of the acceptor ATTO 655 (**Fig. 2c**) as well as the growth of the GUV and expansion of its original basal membrane (white dashed line in **Fig. 2c**). The plots in **Fig. 2d** show the fluorescence intensity of membrane conjugated dyes ATTO 488 and ATTO 655 during liposome delivery. The elevated intensity of ATTO 655 over time confirms fusion and lipid mixing^*12*^ (*cf*. **S1** for calculation of FRET efficiency). The membrane area and the internal volume of the GUV before and after liposomal fusion (**Fig. 2a-c**) are shown in **Fig. 2e**. The basal membrane increases by 27% of its initial area. By taking into account the growth of the basal membrane and assuming that the dome-shaped GUV is approximately a spherical cap with a consistent height measured from z-scan, we determined the total membrane area increase to be 81.22 μm^2^ (*cf*. **S2** for calculation of GUV membrane area). If the diameter of each SUV is assumed to be 100 nm, this area increase corresponds to approximately 2580 fused liposomes. Eventually, the GUV’s volume increases by 15% of its initial volume (*cf*. **S2** for calculation of GUV volume).

**Figure 2.**
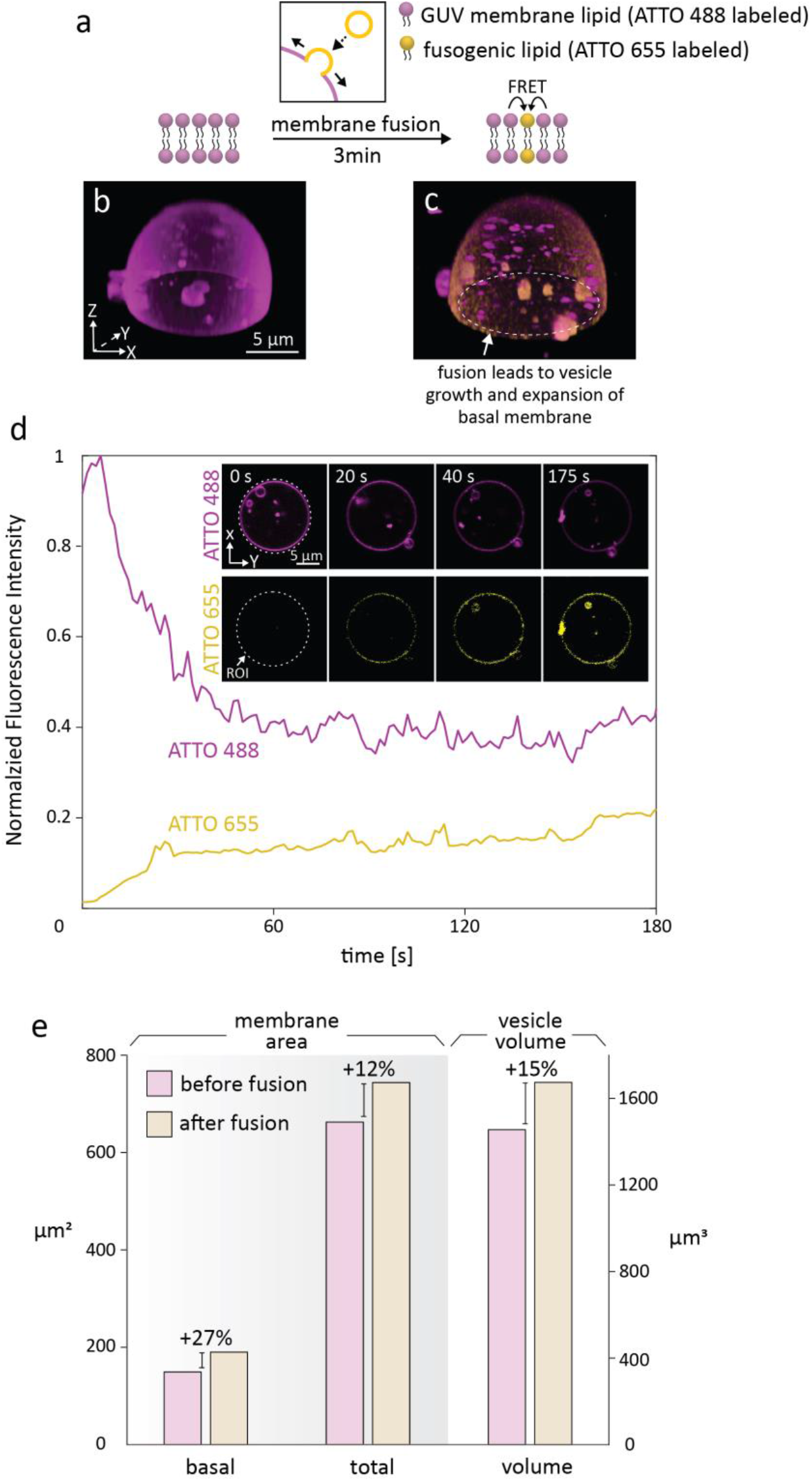
Fusion of giant vesicles with liposomes. (**a-d**) Membrane fusion and mixing was monitored by labeling the membrane of the GUV (magenta) and the liposomes (yellow) with a FRET pair. (**a**) Schematic drawing depicting the membrane fusion visualized by FRET between donor-labeled (ATTO 488) GUV membrane lipid and acceptor-labeled (ATTO 655) fusogenic lipid. (**b-c**) 3D confocal micrograph of a surface-adhered GUV before (**b**), and after (**c**) fusion. Emission channels of ATTO 488 and ATTO 655 are overlaid in the micrograph; only ATTO 488 was excited. The GUV grows and its basal membrane expands upon membrane fusion. (**d**) Plots show that the fluorescence intensity of ATTO 655 (yellow) increases, and ATTO 488 (magenta) decrease over time. (*cf*. Materials and Methods for details of normalization) The inset shows the xy cross-sectional view of the GUV in the ATTO 488 and ATTO 655 channels at 0, 20, 40 and 175 s. The fluorescence intensity is measured in the region of interest (ROI, encircled in dotted line). (**e**) Histograms depicting the changes in GUV basal area, GUV total area and GUV volume before, and after 3 minutes of liposome exposure.

Electrostatic interactions play a dominating role in membrane fusion^*43, 44*^. Lira *et al*. reported that full fusion only occurs between giant vesicles and liposomes if the membranes contain a certain percentage of oppositely charged lipids e.g., minimum 20 mol% anionic lipids in GUV and 48 mol% of cationic lipids in the liposomes^*25*^. In our system, 52 mol% of liposomal lipids are cationic (DOTAP), and GUVs consists of 71 mol% PE (neutral), 23 mol% PG (-) and 5 mol% CA (2-). Full fusion of membrane is required for cargo release^*45*^.

When we observed the spreading of basal membranes during liposome fusion, GUVs occasionally disintegrated and collapsed during this process (*cf*. **S3**). This can be due to an excessive increase in membrane tension as a result of the strong adhesion between the membrane and surface^*46-48*^.

### Delivery of cargo

We successfully encapsulated various cargo in GUVs upon liposomal delivery (**Fig. 3**). Prior to the delivery, the internalization of cargo inside the liposomes was achieved based on the established methods for transfection/lipoplex formation (*cf*. Materials and Methods for details). To illustrate that cargo cannot diffuse into the GUV without liposomal delivery, we exposed RNA oligomer conjugated with fluorescein amidite (FAM) (green color) directly to the GUV (magenta color) (**Fig. 3a-b**). After 120 s of exposure, almost no RNA can be detected in the GUV (**Fig. 3b**). In contrast, RNA delivered in fusogenic liposomes was transported into the GUV within 70 s (**Fig. 3c-d**). Here, both the GUVs and the liposomes were fluorescently labeled with the same fluorophore (ATTO 655-DOPE, false colored in magenta). The confocal microscopy time series of the vesicle depicted in **Fig. 3d** is shown in **Fig. 3e-g**. The basal membrane of the GUV expanded during this period. **Fig. 3h** shows the mean fluorescence intensity of RNA in the region of interest (ROI) indicated with a green dotted line in **Fig. 3e**. The position and size of this circular ROI was unchanged throughout the time of the measurement. Similarly, ssDNA labeled with FAM (**Fig. 3i-j**), pyranine (**Fig. 3k-l**), dextran labeled with fluorescein isothiocyanate (FITC) (**Fig. 3m-n**) and avidin labeled with Alexa Fluor 488 (AF488) (**Fig. 3o-p**) were successfully delivered to the GUV.

**Figure 3.**
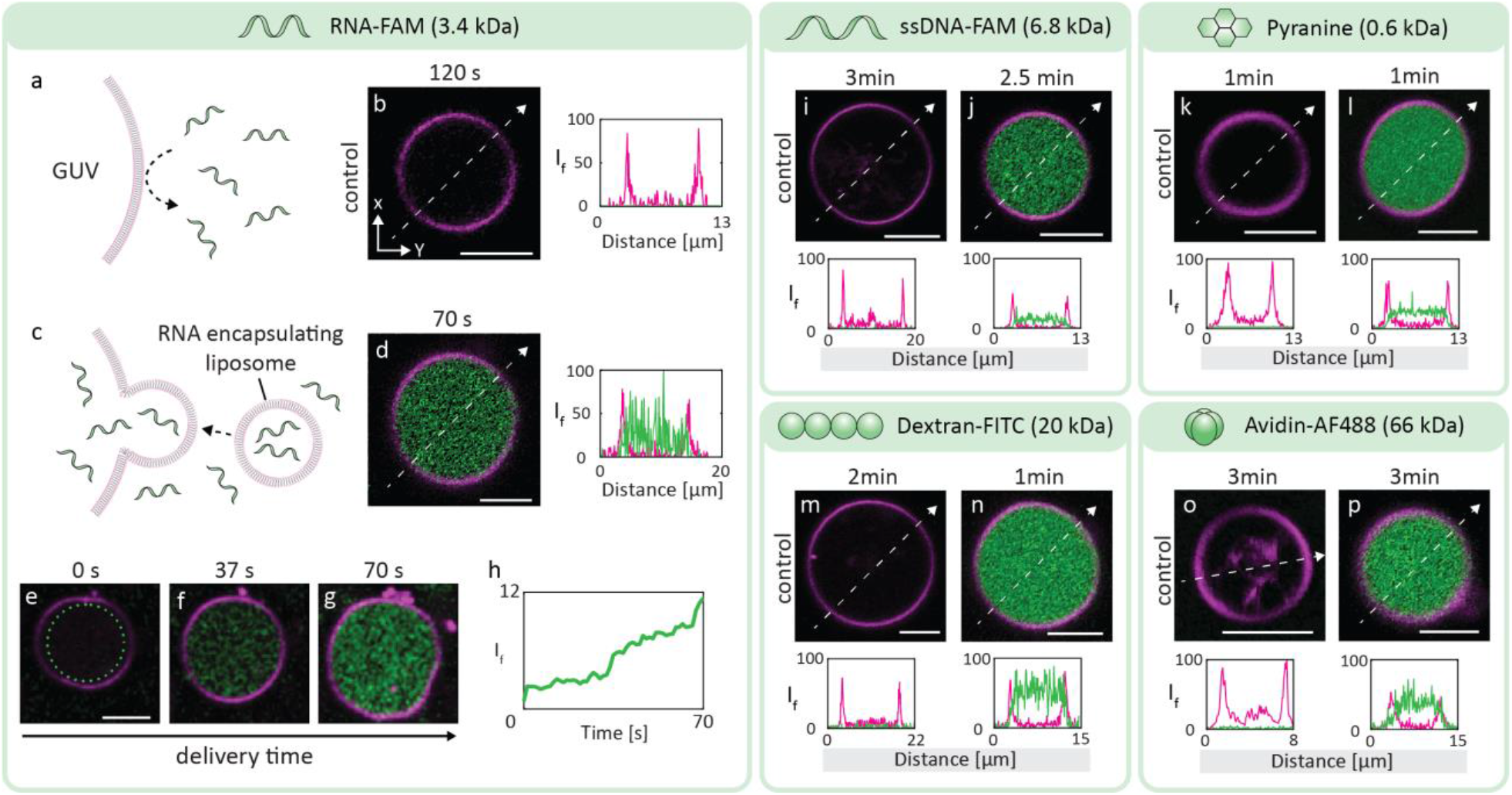
Delivery of membrane-impermeable cargo to GUV. (**a-b**) Fluorescently labeled RNA oligomers (green) did not permeate the GUV membrane (magenta) within 120 s of direct exposure (control). (**c-d**) When RNA was loaded into the fusogenic liposomes, it was delivered to the GUVs within 70 s. (**e-h**) Confocal microscopy time series showing the liposomal delivery of RNA to the GUV shown in (**d**). (**h**) Fluorescence intensity of internalized RNA inside the ROI (green dotted circle) versus time. The same set of experiments was performed for the delivery of ssDNA-FAM (**i-j**), pyranine (**k-l**), dextran-FITC (**m-n**) and avidin-AF488 (**o-p**). All florescence images are false colored, and shown in an xy cross-sectional view. Scale bars: 5 μm.

The control experiments (**Fig. 3b, i, k, m, o**) confirmed that the investigated GUV membranes were not permeable to the cargo molecules we delivered to the giant compartments. Successful delivery of these molecules to the GUVs was only observed via the liposome fusion route.

We show in the supplementary information further attempts to deliver additional cargo, e.g. temperature-responsive polymer Poly(N-isopropylacrylamide) (PNIPAM) (**S4**), and the lipophilic dye Rhodamine 123 (Rho123) (**S5**). In the PNIPAM case, the SUVs were labeled with ATTO 655 in order to observe potential membrane mixing, similar to the experiment depicted in **Fig. 2a**. Mixing did not occur; we were thus not able to encapsulate the polymer (**S4**). It is conceivable that the presence of the polymer outside the SUVs in the delivery suspension hindered efficient contact of the SUVs to the GUV, preventing fusion. Similarly, we exposed free Rhodamine 123 to the GUV (control), and we observed a low degree of encapsulation by the GUV within 1 min of superfusion (**Fig. S5a-b**) followed by rapid leakage out of the container. When a GUV was exposed to Rho123-containing SUVs, the fluorescence intensity of internalized cargo matched the intensity of the background (**Fig. S5c-d**). Similarly, immediate leakage of cargo was observed after suspension of the superfusion. We attribute the rapid leakage to the low molecular weight (344 Da) and low polar surface area (85 Å^2^) of the molecule^*49*^, which enables its penetration through a bilayer membrane by passive diffusion.

### Enzymatic reaction

We delivered the enzyme CA and performed an enzymatic reaction^*31, 50-52*^ inside lipid compartments. In this reaction CA catalyzes the transformation of the membrane permeable, non-fluorescent substrate CFDA into CF, which is membrane impermeable and fluorescent^*31, 50*^. To perform the reaction inside the GUVs, first the enzyme was encapsulated within the liposomes, and subsequently delivered to the GUVs (Step 1 in **Fig. 4a**). Thereafter, the giant compartment was exposed directly to free CFDA, which is able to diffuse across the membrane (Step 2 in **Fig. 4a**). Enzymatic hydrolysis of the substrate inside the GUV led to CF formation. Upon hydrolysis, two acetate groups are cleaved off (green dashed lines on molecular structure of CFDA in **Fig. 4a**), forming polar hydroxy groups which decreased the lipophilicity of the product (CF) by ten-fold, as compared to the substrate (CFDA)^*53*^. The product became membrane impermeable and accumulated inside the GUV, causing a gradual increase in the fluorescence signal (Step 3 in **Fig. 4a**).

**Figure 4.**
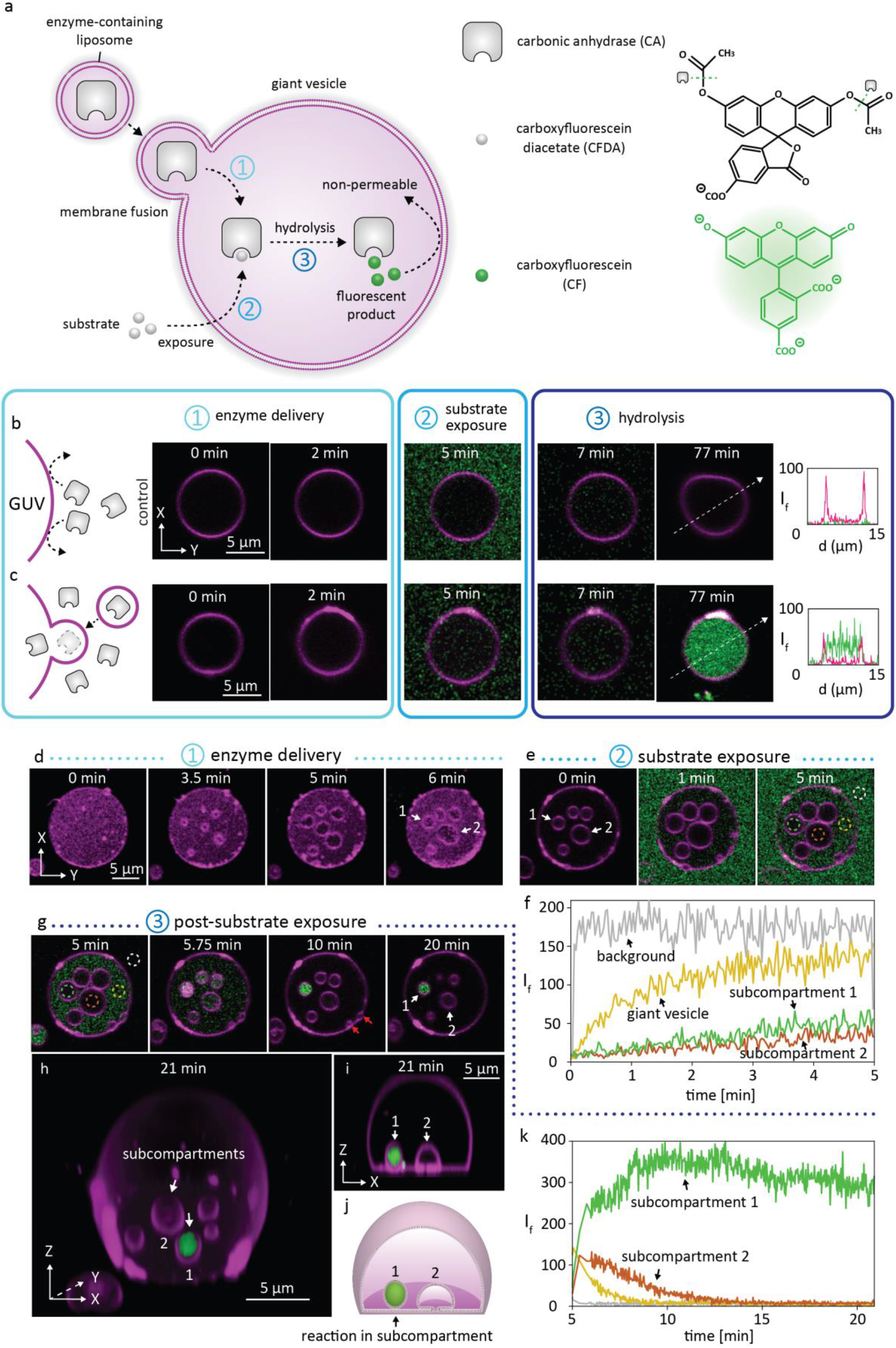
Delivery of an enzyme and observation of enzymatic reaction inside GUVs. (**a**) Schematic drawing summarizing the experimental process. (1) Enzyme (CA) is delivered via liposomes and encapsulated by a GUV. (2) This is followed by the direct delivery and diffusion of non-fluorescent substrate (CFDA) into the GUV. (3) Internalized enzyme catalyzes the hydrolysis of the substrate which leads to production of membrane impermeable fluorescent CF (green colored). (**b**) Direct delivery of the enzyme followed by the substrate exposure. (**c**) Liposomal delivery of the enzyme followed by substrate exposure. Accumulation of CF can be observed inside the lipid compartment. (**d-k**) Enzymatic reaction followed inside a subcompartmentalized GUV. (**d**) Several subcompartments form and grow during the enzyme delivery (two of them are numbered -#1 and #2- and indicated with white arrows). (**e**) GUV is exposed to the substrate. Four ROIs are encircled in dashed lines: background/outside the vesicle (grey), primary volume of the giant vesicle (yellow), subcompartment 1 (green) and subcompartment 2 (orange). (**f**) Plots showing the fluorescence intensity measured in each ROI in (**e**), versus time. (**g-k**) CF formation in subcompartments after substrate was exposed, over time. Red arrows in (**g)** indicate small vesicles adhered to the protocell membrane. (**h-i**) Confocal micrograph of the subcompartmentalized GUV in 3D (xyz) (**h**) and in side view (xz cross-sectional) (**i**). All other confocal micrographs are shown in top down (xy cross-sectional) view. (**j**) Schematic drawing corresponding to (**i**). (**k**) Plots showing the fluorescence intensity of CF in each ROI in (**g**) over time.

**Fig. 4b** shows the control experiment, where a giant vesicle was first exposed to CA for 2 min directly, and subsequently to CFDA for 5 min. Fluorescence was not detectable inside the GUV after 77 min, indicating that no enzyme was internalized, and no CF could be produced.

In contrast, we observed a measurable increase in fluorescence intensity inside the giant vesicle shown in **Fig. 4c** over the course of 77 min, where the CA was delivered with SUVs before substrate exposure. This confirms successful internalization of the enzyme. We note that the experiments were performed at room temperature rather than the optimal 37 °C, which is the likely reason for the relatively long reaction time observed for CF generation.

During the enzyme delivery to another GUV, we observed formation of several membranous subcompartments (**Fig. 4d**). We recently reported the unusual compartment formation on the base of surface-adhered GUVs^*4*^, which could be, in the context of the origins of life, a conceivable way of division and generation of functional diversity within primitive containers. After enzyme delivery to the GUV, the subcompartmentalized vesicle was exposed to the substrate (**Fig. 4e**) and the fluorescence intensities of several ROIs shown in **Fig. 4e** (encircled in dashed lines), were monitored and plotted over time (**Fig. 4f**).

We observe weak fluorescence of the substrate prior to its conversion into CF (**Fig. 4b, 4c** and **4e**). This is due to an impurity in the substrate, which is present due to unavoidable non-enzymatic hydrolysis of the carboxyfluorescein diacetate^*54, 55*^.

The plots in **Fig. 4f** reveal increasing fluorescence inside the primary volume of the giant compartment over time, and at a lower magnitude in the subcompartments. Subsequently, we terminated the exposure of the substrate to the GUV and continuously monitored the fluorescence emission. The fluorescence intensities of the primary volume, as well as of the subcompartment #2, gradually decreased (**Fig. 4g**), indicating that most of the substrate (CFDA) leaked out before it could be converted to the product (CF). Since the subcompartmentalization is a result of tension-driven membrane flow from areas of high to areas of low membrane curvature, we anticipate that more transient pores^*4, 56*^ open in this particular compartment, which is the likely the cause of leakage of the substrate. Disappearance of small vesicles at the protocell membrane (red arrows in **Fig. 4g**), indicating rapid membrane remodeling, support our hypothesis. The fluorescence intensity inside the subcompartment #1, however, rapidly increased (**Fig. 4g-k**), reaching a peak intensity at around 10 min (green line in **Fig. 4k**). The slow decrease of intensity afterwards might be due to photobleaching of CF during the prolonged imaging (**Fig. 4g**). **Fig. 4h** shows a confocal micrograph of the giant vesicle in 3D, and **Fig. 4i**, the xz cross-sectional view at 21 min. Unlike subcompartment #2, #1 appears to be completely sealed or connected to the basal membrane through a nano-sized tether, which would not be visible in the micrograph^*4*^ (**Fig. 4i-j)**. #2 is semi-grown with a visible conduit to the surface (**Fig. 4i-j)**. In this case, while the contents of subcompartment #2 can escape, the reactants and products inside subcompartment #1 remain confined.

It appears that the enzymatic reaction in subcompartment #1 is amplified. We attribute this to a direct entry path into the subcompartment via the gap between the basal membrane and the surface, which allows entry into at least some, perhaps even all subcompartments before some of them seal. Since the enzyme-encapsulating liposome suspension was not purified (e.g. by dialysis, gel filtration or centrifugation), free enzyme from the surrounding solution can be internalized through the gap between the surface and the basal membrane^*4*^. Alternatively, active transport via Marangoni flow can transport membrane-adsorbed free cargo (e.g. enzyme) to the subcompartments^*4, 57*^. In this case, we can assume that all subcompartments should have a similar amount of enzyme internalized. However, even though much enzyme was present in this case, only the rapidly sealing compartments, such as #1, would retain the fluorescent reaction product. Multicompartmentalized giant vesicles have also been shown to accommodate multi-step reactions based on the diffusive translocation of reactants through a bilayer, mediated by protein-stabilized membrane pores^*7, 58, 59*^. In our subcompartmentalized reaction system, the involvement of transmembrane proteins is not necessary for efficient uptake of large molecules or chemical reactants.

## Conclusion

In this work, we showed the internalization of cargo in selected surface-adhered GUVs by efficient membrane fusion between small cationic liposomes and the giant vesicles. Fusion of approximately 2580 small unilamellar vesicles results in a volume growth of 15% in a giant vesicular compartment within a few minutes. Upon vesicle fusion, small molecules as well as biopolymers were transferred to, and retained inside surface-adhered giant vesicles. An enzymatic reaction was also demonstrated inside a GUV and its subcompartment following internalization of the catalyst. The integration of this enzymatic reaction inside a GUV subcompartment not only confirms earlier findings on chemical communication to subcompartments, but also shows that a mechanism exists that can selectively encapsulate cargo in individual subcompartments. Such a process may explain the development of diversified functionality in organelle-like units inside synthetic cells, or substructured protocells in the context of the origins of life.

The experimental technique described here is potentially useful and convenient for implementing pre-or probiotic reactions and reaction networks in model amphiphilic protocells^*60*^, and cell-free gene expression models in surface-adhered artificial cells.

## Material and Methods

### Preparation of giant vesicle suspension

The dehydration and rehydration method^*16*^ were used to prepare the lipid suspensions. The following lipids and lipid-conjugated fluorophores were used: 1,2-Dioleoyl-3-trimethylammonium-propane (DOTAP; Avanti Polar Lipids, USA), 1,2-Dioleoyl-sn-glycero-3-phosphoethanolamine (DOPE; Avanti Polar Lipids, USA), E. coli L-α-phosphatidylethanolamine (E. coli PE; Avanti Polar Lipids, USA), E. coli L-α-phosphatidylglycerol (E. coli PG; Avanti Polar Lipids, USA), E. coli Cardiolipin (E. coli CA, Avanti Polar Lipids, USA), ATTO 488- and ATTO 655-conjugated DOPE (Atto-Tech GmbH, Germany). Lipid membranes with two different lipid compositions were prepared: DOTAP:DOPE:ATTO 655-DOPE (w/w: 50:49:1) to form liposomes and PE:PG:CA:ATTO 488-DOPE (or ATTO 655-DOPE) (w/w: 67:23:9:1) to form GUVs.

A total of 3000 µg of lipids in 300 μL of chloroform (10 μg/μL) was placed in a 10 mL round bottom flask. The chloroform was removed in a rotary evaporator at reduced pressure (20 kPa) for 6 hours. The dry lipid film was rehydrated with 3 mL of a phosphate-buffered saline (PBS) buffer containing 5 mM Trisma base, 30 mM K_3_PO_4_, 30 mM KH_2_PO_4_, 3 mM MgSO_4_·7H_2_O and 0.5 mM Na_2_EDTA (pH=7.4, adjusted with 3 M H_3_PO_4_), and 30 μL of glycerol, and kept at 4 °C for overnight. The next day, the rehydrated lipid film was sonicated in a water bath with an ultrasonic cleaner (USC-TH, VWR) at 20 °C for 10-20 s to obtain giant vesicle suspension (uni- and multilamellar).

Unless specified otherwise all chemicals were purchased from Sigma-Aldrich (USA).

### Surface fabrication and preparation of the observation chamber

SiO_2_ surfaces were fabricated at the Norwegian Micro- and Nano-Fabrication Facility at the University of Oslo (MiNaLab). Thin films (≈84 nm) were deposited on circular borosilicate glass cover slips (#1.5H, Menzel Gläser) by E-beam physical vapor deposition (EvoVac, Ångstrom Engineering, Canada).

The observation chamber was prepared by gently adhering a Poly(dimethyl siloxane) (PDMS) frame onto SiO_2_ surfaces and filling the frame with 4 mL of HEPES buffer containing 4 mM CaCl_2_.

### Preparation of surface-adhered GUVs

4 μL of the giant vesicle lipid suspension described above with the specific composition PE:PG:CA:ATTO 488/655-DOPE was desiccated for 20 min and rehydrated with 200 μL of HEPES buffer containing 10 mM HEPES and 100 mM NaCl (pH=7.8, adjusted with 5 M NaOH) for 10 min and transferred to the observation chamber via an automatic pipette.

### Preparation of cargo-free liposomes

Liposomes composed of DOTAP:DOPE:ATTO 655-DOPE were prepared by extrusion on the same day of the experiments. First, 50 μL of giant vesicle lipid suspension was diluted with Tris buffer containing 125 mM NaCl, 10 mM Trisma base and 1 mM Na_2_EDTA, (pH=7.4, adjusted with 3 M HCl) to a total lipid concentration of 0.1 mg/ml. The diluted suspension was sonicated for 5 min in an ultrasonic cleaner at 20°C, and extruded through a 100 nm pore size filter 11 times by using a mini extruder (Avanti Polar Lipids, USA). The extruded sample was stored at 4°C until use.

### Preparation of RNA-, DNA- and avidin-containing liposomes

For the preparation of liposomes encapsulating 10-base RNA oligomers (5′-FAM-AAA AAA AAA A-3′, 3.4 kDa, Dharmacon, USA), 20-base ssDNA (5’/56-FAM/TGT ACG TCA CAA CTA CCC CC-3’, 6.8 kDa, Integrated DNA Technologies, USA) and avidin-Alexa Fluor 488 (66 kDa, Invitrogen, USA), the protocol leading to cargo-free liposomes was used with a slight modification. The dry lipid film to form the giant vesicle suspension was rehydrated with ethanol (99.9%, Antibac AS, Norway) instead of PBS buffer, based on a previously established protocol^*61*^. Briefly, the dry lipid film was rehydrated with 600 μL of ethanol to reach a lipid concentration of 5 μg/μL, followed by sonication in an ultrasonic cleaner for 2 min at 40 °C.

To obtain a lipid-cargo suspension, each type of cargo molecule specified above was dissolved in Tris buffer and mixed with the ethanol-lipid solution resulting in a weight ratio of 10:1, lipid: cargo molecule. The volume of ethanol-lipid solution added to the cargo solution was kept at a maximum of 10% of the lipid-cargo suspension volume. The lipid-cargo suspension was then sonicated for 5 min and extruded 11 times through a 100 nm pore size filter. The final concentration of each encapsulated component was 8 μM for RNA, 7.68 μM for DNA and 2 μM for avidin, and the final lipid concentration was 0.1 μg/μL.

### Preparation of pyranine-and dextran-containing liposomes

For the preparation of liposomes containing pyranine (8-Hydroxypyrene-1,3,6-trisulfonic acid trisodium salt, 0.6 kDa) and dextran-FITC (20 kDa), 900 μL of Tris buffer containing cargo (100 μM) was mixed with 100 μL of giant vesicle suspension (DOTAP:DOPE:ATTO 655-DOPE). The lipid-cargo suspension was then sonicated for 5 min and extruded 11 times through a 100 nm pore size filter. The concentration of each encapsulated component in the final suspension was 90 μM, and the final lipid concentration was 0.1 μg/μL.

### Preparation of liposome-free cargo solutions (control)

For RNA, DNA and avidin, the same protocol to prepare liposomes carrying these components was followed, except that the ethanol-lipid solution was substituted by the same volume of ethanol which did not contain any lipid.

For pyranine and dextran, the same protocol to prepare the liposomes carrying these components was followed, except that 900 μL of cargo solution in Tris buffer was mixed with 100 μL of PBS buffer as a substitute for the lipid-cargo suspension.

### Exposure of surface-adhered GUVs to fusogenic liposomes or liposome-free cargo

Fusogenic liposomes with or without cargo, as well as liposome-free cargo solutions were locally exposed to surface-adhered GUVs using an open-space microfluidic pipette (Biopen, Fluicell AB, Sweden). Briefly, the microfluidic pipette was positioned 10-20 μm above the surface using a 3-axis water hydraulic micromanipulator (Narishige, Japan), and by activating the device, an exposure zone containing liposomes around the surface-adhered GUV isolated from the ambient buffer was created.

### Enzymatic reaction in GUVs

Tris buffer containing 5 mM Trisma base (PH=7.5, adjusted with 3 M HCl) was used to dissolve CA (Carbonic Anhydrase I from human erythrocytes, 30 kDa)^*31*^. To form CA-encapsulating liposomes, the same protocol used for RNA/DNA/avidin was employed. The final ethanol-TRIS buffer mixture contained 2 μM CA and 0.1 μg/μL of lipids.

For the preparation of the free CA solution (control), the same protocol was followed except that the ethanol-lipid solution was substituted by same volume of ethanol free of lipids.

CFDA (6-Carboxyfluorescein diacetate, ∼95%, 0.46 kDa) solution was prepared by adding 3 μL of 5 mM CFDA in dimethyl sulfoxide (DMSO) to 97 μL of Tris buffer containing 5 mM Trisma base (PH=7.5, adjusted with 3 M HCl), to achieve a CFDA concentration of 150 μM^*31*^.

CA-encapsulating liposomes, liposome-free CA (control) and CFDA were exposed to GUVs by using the microfluidic pipette.

### Microscopy and image analysis

For imaging, a laser scanning confocal microscopy (Leica SP8) equipped with a HCX PL APO CS 63x oil objective (NA 1.4) was used. The excitation/emission wavelengths for the fluorophores were: 458/510-600 nm for pyranine, 500/510-600 nm for ATTO 488, Alexa Fluor 488, FAM, FITC, and 660/680-790 nm for ATTO 655. The 3D fluorescence micrographs in **Fig. 2b, c** and **Fig. 4h** were reconstructed using the Leica Application Suite X Software (Leica Microsystems, Germany). The intensity analyses of the micrographs in **Fig. 2d, 3b-p, 4b-c, e-g, k** were also performed using the Leica Application Suite X Software. Schematic drawings were created with Adobe Illustrator CS6 (Adobe Systems, USA).

In **Fig. 2d**, the fluorescence intensities of both ATTO 488 and ATTO 655 are normalized by the maximum intensity of ATTO 488 during 3 min of fusion 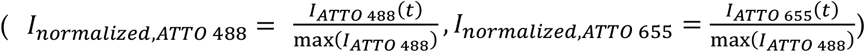. All graphs in **Fig. 2, 3, 4, S1, S5** were plotted using MATLAB R2020b.

## Acknowledgements

This work was made possible through financial support obtained from the UiO: Life Sciences Convergence Environment, the Research Council of Norway (ForskningsrÅdet) Project Grant 274433, as well as the startup funding provided by the Centre for Molecular Medicine Norway (RCN 187615), and the Faculty of Mathematics and Natural Sciences at the University of Oslo.

A.B.S. gratefully acknowledges financial support of the National Science Foundation Graduate Research Fellowship (NSF GRFP) – Graduate Research Opportunities Worldwide (GROW) under grant No. DGE1745303.

