## Supporting Information for "Liposome-assisted in-situ cargo delivery to artificial cells and cellular subcompartments"

### S1. Calculation of FRET efficiency

FRET efficiency ($E_{FRET}$) was determined as: $E_{FRET} = I_{ATTO\,655} / (I_{ATTO\,488} + I_{ATTO\,655})$[1, 2]. $I_{ATTO\,488}$ and $I_{ATTO\,655}$ are the fluorescence intensities of ATTO 488 ($E_m$=510-600 nm) and ATTO 655 ($E_m$=680-790 nm) respectively, when only ATTO 488 is excited at 500 nm. **Fig. S1** shows the FRET efficiency during 3 min fusion of giant vesicles and fusogenic liposomes.

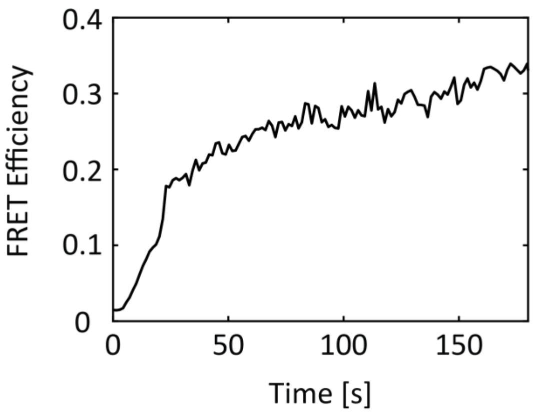

**Figure S1**. FRET efficiency during fusion of giant vesicles and fusogenic liposomes.

### S2. Calculation of GUV membrane area and volume

We assume a surface-adhered GUV is approximately a combination of a spherical cap and a basal membrane (**Fig. S2**). The radius of the basal membrane before and after the fusion is denoted as $r_0$ and $r_t$, respectively. The growth of the basal membrane radius is $\Delta r = r_t - r_0 = 7.775\ \mu m - 6.894\ \mu m = 0.881\ \mu m$. The height of the spherical cap, $h$, measured from z-scan in confocal micrograph, remains approximately the same before and after the fusion: $h_0 = h_t = h = 10.760\ \mu m$. The increase of area of basal membrane is given as $\Delta A_{basal} = A_{basal,t} - A_{basal,0} = \pi r_t^2 - \pi r_0^2 = 40.610\ \mu m^2$ and the spherical cap $\Delta A_{cap} = A_{cap,t} - A_{cap,0} = \pi(r_t^2 + h^2) - \pi(r_0^2 + h^2) = \pi(r_t^2 - r_0^2) = 40.610\ \mu m^2$.

The increase in total membrane area of GUV is calculated as $\Delta A = \Delta A_{basal} + \Delta A_{cap} = 40.610\ \mu m^2 + 40.610\ \mu m^2 = 81.220\ \mu m^2$. The increase of GUV volume is determined as $\Delta V = V_t - V_0 = \frac{1}{6}\pi h(3r_t^2 + h^2) - \frac{1}{6}\pi h(3r_0^2 + h^2) = \frac{\pi h(r_t^2 - r_0^2)}{2} = 218.480\ \mu m^3$.

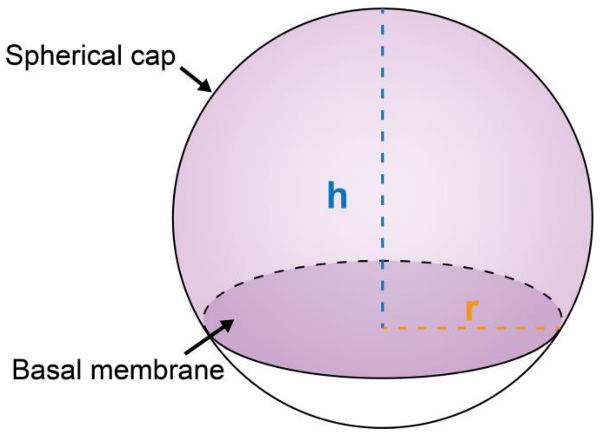

**Figure S2**. Schematic drawing of a GUV approximated as a spherical cap with a basal membrane.

### S3. Disintegration of GUV during the membrane fusion

**Fig. S3** shows two occasions of disintegration of surface-adhered GUV upon membrane fusion. In **Fig. S3a,b**, the GUV partially spread on surface after 4 min of membrane fusion. In **Fig. S3c,d**, the GUV disintegrated and collapsed after 2 min of membrane fusion.

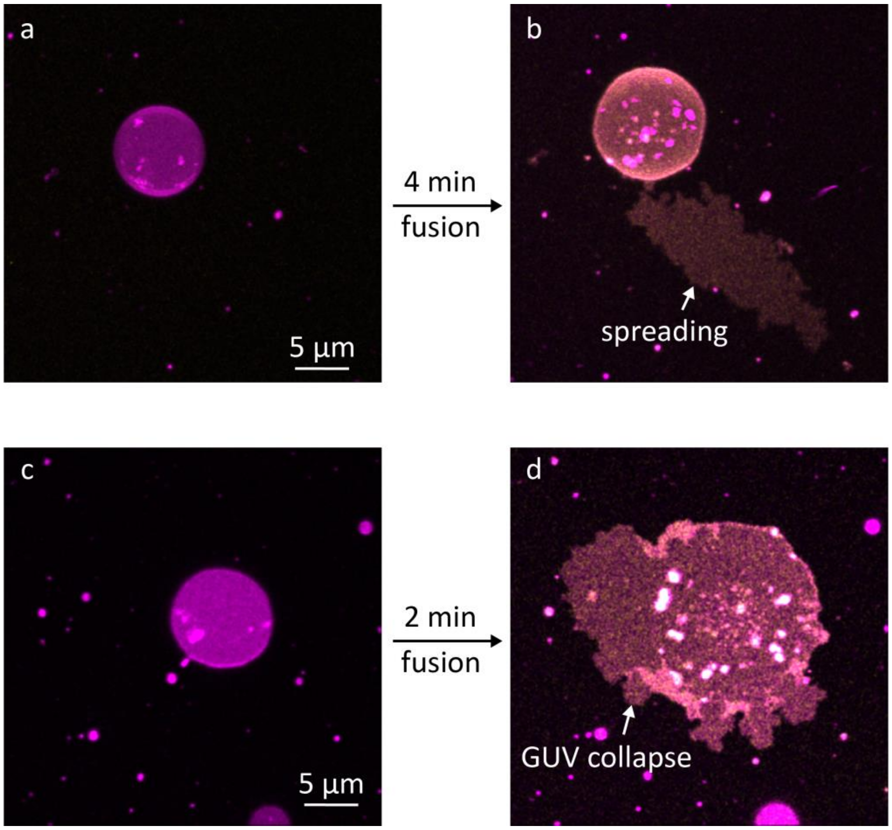

**Figure S3**. Spreading and collapse of GUV upon membrane fusion from top view. (**a,c**) Surface-adhered GUV labeled with ATTO 488 (magenta) was exposed to fusogenic liposomes labeled with ATTO 655 (yellow) resulting in partial spreading (**b**) or collapse of GUV (**d**) on the surface. ATTO 488 and ATTO 655 channels are overlapped and the base of GUV is in focus.

### S4. Delivery of PNIPAM

Liposomes containing PNIPAM (30 kDa) were prepared by mixing 900 µL of cargo solution in Tris buffer (60 mg/mL) with 100 µL of giant vesicle suspension (DOTAP:DOPE:ATTO 655-DOPE). The lipid-cargo suspension was then sonicated for 5 min and extruded 11 times through a 100 nm pore size filter. The final concentration of PNIPAM was 54 mg/mL and the final lipid concentration was 0.1 µg/µL.

The lipid-PNIPAM suspension was exposed to GUVs by using the microfluidic pipette. We examined the delivery of PNIPAM to GUV by monitoring the membrane fusion of GUV and PNIPAM-encapsulating liposomes. GUV (magenta color) was exposed to PNIPAM-encapsulating liposomes (yellow color, **Fig. S4a-c**), where the membranes of the GUV and the liposomes were labeled with ATTO 488 and ATTO 655, respectively, and both fluorophores were excited. After 2.5 min of exposure, no increase in fluorescence signal in ATTO 655 channel (**Fig. S4e)** thus no fusion occurs, and therefore we conclude that the delivery of PNIPAM to GUV is not successful.

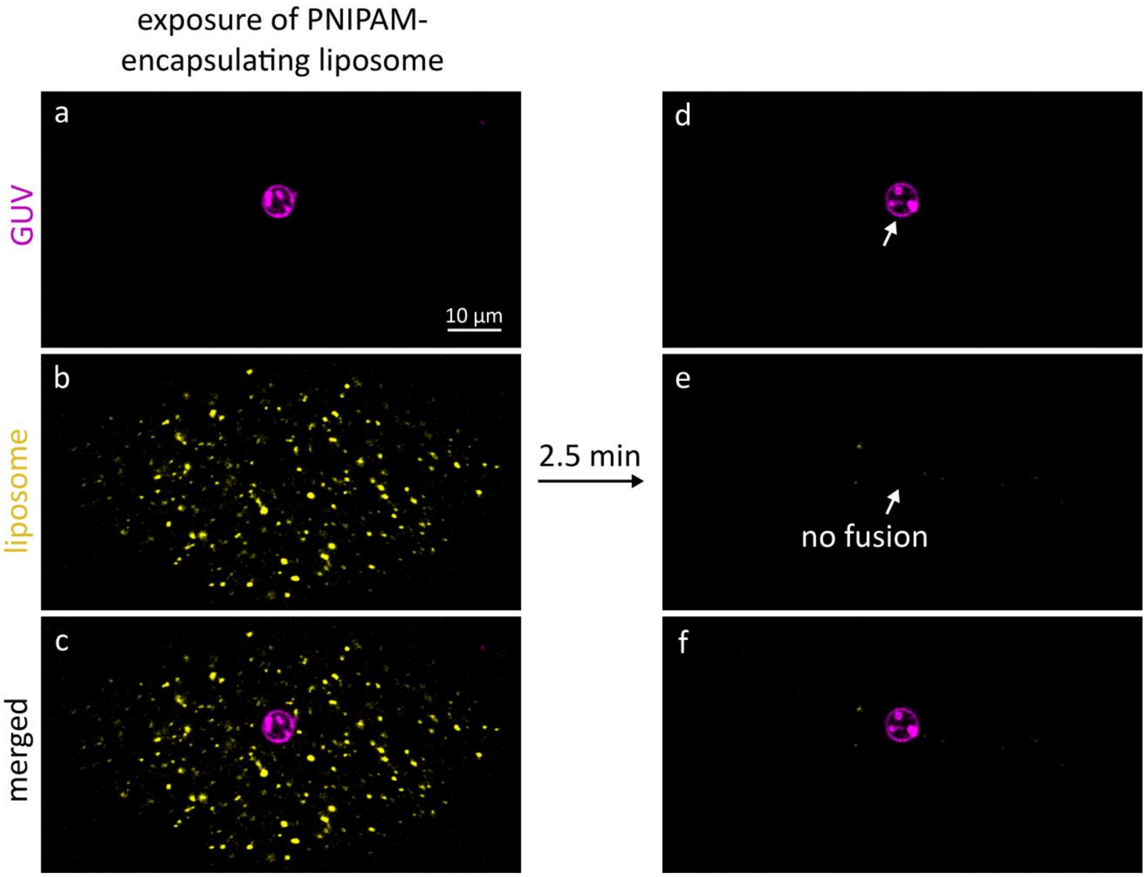

**Figure S4**. Exposure of PNIPAM-encapsulating liposomes to a GUV. (**a**,**d**) shows the fluorescence of GUV, and (**b**,**e**) the fluorescence of fusogenic liposome. (**c**) and (**f**) shows the (**a**-**b**) and (**d**-**e**) overlaid, respectively.

#### S5. Delivery of Rhodamine 123

Liposomes containing Rhodamine 123 (≥85%, 0.38 kDa) was prepared by rehydrating the dry lipid film (DOTAP:DOPE:ATTO 655-DOPE) described in Materials and Methods with 3 ml of 500 µM Rhodamine 123 in HEPES buffer was used to, followed by 1 min of vortexing and 10 min of sonication in an ultrasonic cleaner at 20 °C. The suspension was diluted 10 times in HEPES buffer to achieve a Rhodamine 123 concentration of 50 µM and a lipid concentration of 0.1 µg/µL, and was sonicated in an ultrasonic cleaner at 20 °C for 5 min, finalized by 11 times of extrusions through a 100 nm pore size filter.

For the control experiment, the protocol mentioned above was followed, except that in the step of rehydration of the dry lipid film, Rhodamine 123 in HEPES buffer without lipid was subjected to vortexing and sonication to obtain the suspension.

We performed the delivery of Rhodamine 123 by exposing free Rhodamine 123 (**Fig. S5a-b**) or Rhodamine 123-encapsulating liposomes (**Fig. S5c-d**) to a GUV by using the microfluidic pipette. In control experiments (**Fig. S5b**), the internalized Rhodamine 123 (green plots) intensity appears to be much less than in the background (grey plots). When the GUV was exposed to Rhodamine 123-encapsulating liposomes **(Fig. S5d)**, the internalized Rhodamine 123 (green plots) intensity follows the same trend as the background fluorescence (grey plots), indicating an extremely high delivery efficiency. However, in both cases, the encapsulated Rhodamine 123 leaked immediately once the exposure was stopped.

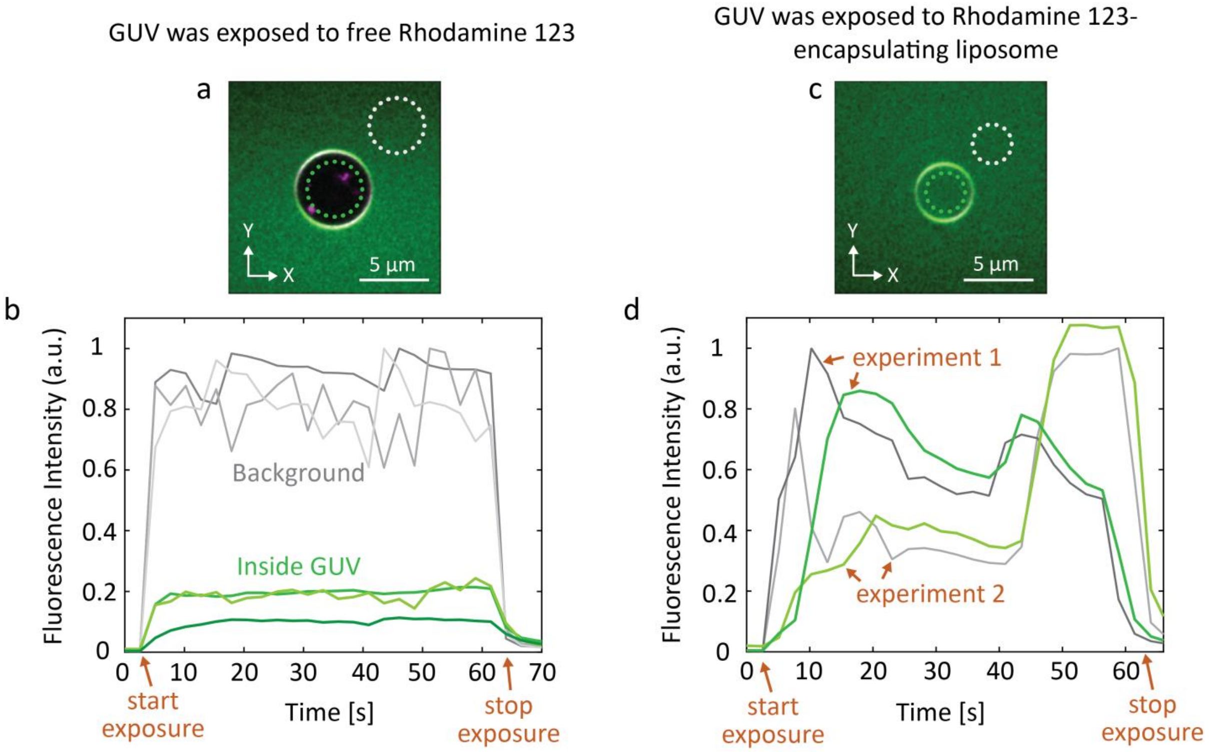

**Figure S5**. Delivery of Rhodamine 123 to GUV. (**a**-**b**) GUV was exposed to free Rhodamine 123 (control). (**c**-**d**) GUV was exposed to Rhodamine 123-encapsulating liposomes. (**a**,**c**) Rhodamine 123 fluorescence (green) is overlaid with GUV fluorescence (magenta). (**b**,**d**) Plots in green color represent Rhodamine 123 fluorescence inside GUV (encircled in green dash line in **a** and **c**) over time and plots in gray color, the background fluorescence (encircled in grey dash line in **a** and **c**). (**b**) includes 3 individual experiments, and (**d**), 2 individual experiments.
